# Identification and Ranking of Recurrent Neo-Epitopes in Cancer

**DOI:** 10.1101/389437

**Authors:** Eric Blanc, Manuel Holtgrewe, Arunraj Dhamodaran, Clemens Messerschmidt, Gerald Willimsky, Thomas Blankenstein, Dieter Beule

**Affiliations:** Core Unit Bioinformatics, Berlin Institute of Health, Berlin, Germany; Institute of Immunology, Charité - Universitätsmedizin Berlin, corporate member of Freie Universität Berlin, Humboldt-Universität zu Berlin, and Berlin Institute of Health, Berlin, Germany; Max Delbrück Center for Molecular Medicine in the Helmholtz Association (MDC), Berlin, Germany; Berlin Institute of Health, Berlin, Germany; Charité - Universitätsmedizin Berlin, corporate member of Freie Universität Berlin, Humboldt-Universität zu Berlin, and Berlin Institute of Health, Berlin, Germany; German Cancer Research Center (DKFZ), Heidelberg, Germany

**Keywords:** cancer, immunotherapy, neo-epitope, neo-antigen, precision treatment

## Abstract

Immune escape is one of the hallmarks of cancer and several new treatment approaches attempt to modulate and restore the immune system’s capability to target cancer cells. At the heart of the immune recognition process lies antigen presentation from somatic mutations. These neo-epitopes are emerging as attractive targets for cancer immunotherapy and new strategies for rapid identification of relevant candidates have become a priority. We carefully screen TCGA data sets for recurrent somatic amino acid exchanges and apply MHC class I binding predictions. We propose a method for *in silico* selection and prioritization of candidates which have a high potential for neo-antigen generation and are likely to appear in multiple patients. While the percentage of patients carrying a specific neo-epitope and HLA-type combination is relatively small, the sheer number of new patients leads to surprisingly high reoccurence numbers. We identify 769 epitopes which are expected to occur in 77629 patients per year. While our candidate list will definitely contain false positives, the results provide an objective order for wet-lab testing of reusable neo-epitopes. Thus recurrent neo-epitopes may be suitable to supplement existing personalized T cell treatment approaches with precision treatment options.

## Introduction

Increasing evidence suggests that clinical efficacy of cancer immunotherapy is driven by T cell reactivity against neo-antigens [1], [2], [3], [4] and [5]. While not yet fully understood, immune response and recognition of tumor cells containing specific peptides depends critically on the ability of the MHC class I complexes to bind to the peptide in order to present it to a T cell. Neo-antigens can be created by a multitude of processes like aberrant expression of genes normally restricted to immuno-privileged tissues, viral etiology or by tumor specific DNA alterations that result in the formation of novel protein sequences. Furthermore there is now evidence for neo-epitopes generated from alternative splicing [6] and alterations in non-coding regions [7].

With the advent of affordable short read sequencing, comprehensive neo-antigen screening based on whole exome sequencing has become feasible and many cancer immune therapeutic approaches try to utilize detailed understanding of the neoepitope spectrum to create additional or boost pre-existing T cell reactivity for therapeutic purposes [8], [9]. However, in practice the selection and validation of the most promising neo-epitope candidates is a difficult and time-consuming task. The typical approach is based on the private mutational catalogue of the individual patient: exome sequencing data is subjected to bioinformatics analysis and used to predict neo-epitopes and their binding affinities to the MHC class I complex. Our study aims to complement this approach by a precision medicine perspective. We search and prioritize neo-epitope candidates which have a high potential for neoantigen generation and are likely to appear in multiple patients. These neo-antigens hold the potential for development of *off the shelf T cell therapies* for sub groups of cancer patients. We use epidemiological data to give rough estimates for the expected number of patients in these groups.

Candidate prediction always relies on somatic variant detection workflows and affinity prediction algorithms based on machine learning, see e.g. [10]. Binding prediction far from perfect [11] especially for rarer HLA types, and may also depend on mutational context [12]. Catalogues of the neo-epitope landscape across various cancer entities have been created by various authors [13], [14], [15]. While neoantigen landscape is diverse and sparse [13], here we provide an unbiased, comprehensive ranking of candidates, defined as neo-epitopes arising from recurrent mutations, predicted to be binding to a specific HLA-1 allele. The candidates are ranked according to the expected number of target patients.

## Results

### Recurrent variants and candidates

From the GDC repository [16], we have collected somatic variants for 33 TGCA studies. After removing patients without clinical meta-data, and studies with less than 100 patients, we have selected 1,384,531 high-confidence missense SNPs from 9,641 patients, see methods for details. Using this data, 1,055 variants are deemed recurrent (supplementary table 1), as they can be found in more than 1% of the patients in the respective study cohort. These recurrent variants correspond to 869 unique protein changes, as some appear in multiple cancer entities. 77 of the recurrent variants occur in at least 3% of their cohort (43 unique protein changes).

From these 869 unique protein changes, we have generated candidates that are predicted to be strong MHC class I binders in frequent HLA-1 types that we considered for initial selection. 415 (48%) of them lead to a strong binder prediction. In total, there are 772 candidates that are recurrent in a cancer entity cohort, and predicted as binding for a considered HLA-1 type. These candidates are non-redundant among all the 9-, 10- & 11-mers containing the variant: the selection process retains only the peptide sequence with the lowest predicted IC_50_. Figure 1 and table 1 provide an overview of the variant selection and neo-epitope candidates generation processes, while supplementary table 2 lists all neo-epitopes (weak and strong predicted binders) after removing redundancy.

**Figure 1:**
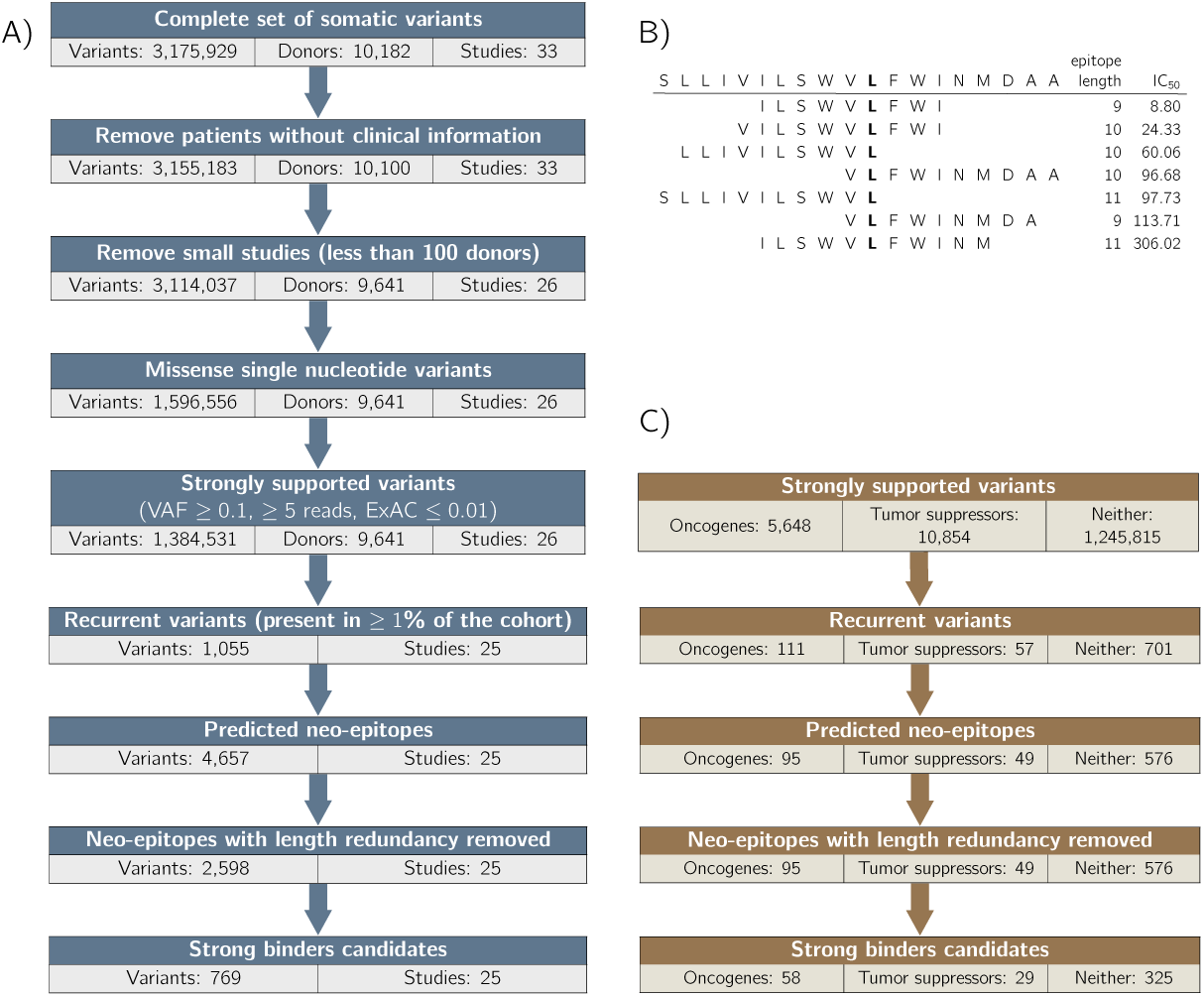
(A) Overview of the recurrent neo-epitope candidates generation process: TCGA studies are selected for at least 100 donors with clinical annotations. For each of these studies, recurrent strongly supported missense Single-Nucleotide Variants are collected. Neo-epitopes binding to 11 HLA-1 types are predicted, redundancy is removed from that set (see B) and strong binders are retained. (B) Example of epitope redundancy: the 18 amino-acids long sequence surrounding recurrent variant GLRA3:S2174L generates 7 binding neo-epitopes for the type HLA-A*02:01. Our pipeline retains only the strongest predicted binder for a given variant and HLA-1 type pair (the first, with an IC_50_ of 8.8 nM in the example). (C) Number of SNVs occuring in genes classified as Oncogenes or Tumor Suppressors by Vogelstein et al. [17], at various point of the variant selection and neo-epitope selection process.

**Table 1:** Overview of the 33 TCGA studies used in this analysis. The 7 studies displayed at the bottom have not been used for the determination of recurrent vairants, as the number of patients is less than 100. The number of strong binders includes all occurrences of neo-epitopes candidates, so a candidate may be counted multiple times when it is predicted to be binding several HLA-1 types.

| Project name | Number of patients |  | Variants per patients |  | Missense variant per patient |  | Recurrent variants | Strong binders |
| --- | --- | --- | --- | --- | --- | --- | --- | --- |
|  | Total | With clinical data | Average | Median | Average | Median |  |  |
| TCGA-BLCA | 412 | 412 | 326 | 226 | 157 | 109 | 22 | 14 |
| TCGA-BRCA | 986 | 986 | 123 | 62 | 50 | 25 | 8 | 10 |
| TCGA-CESC | 289 | 289 | 358 | 157 | 143 | 62 | 17 | 16 |
| TCGA-COAD | 399 | 397 | 666 | 176 | 288 | 82 | 41 | 34 |
| TCGA-ESCA | 184 | 184 | 246 | 187 | 95 | 73 | 80 | 72 |
| TCGA-GBM | 393 | 390 | 212 | 70 | 93 | 36 | 15 | 5 |
| TCGA-HNSC | 508 | 508 | 201 | 139 | 97 | 66 | 14 | 3 |
| TCGA-KIRC | 336 | 336 | 79 | 69 | 33 | 31 | 0 | 0 |
| TCGA-KIRP | 281 | 281 | 85 | 82 | 39 | 38 | 5 | 2 |
| TCGA-LAML | 143 | 143 | 69 | 15 | 16 | 6 | 14 | 7 |
| TCGA-LGG | 508 | 507 | 70 | 36 | 33 | 16 | 14 | 0 |
| TCGA-LIHC | 364 | 364 | 149 | 120 | 70 | 58 | 11 | 16 |
| TCGA-LUAD | 567 | 515 | 367 | 242 | 180 | 113 | 7 | 0 |
| TCGA-LUSC | 492 | 492 | 368 | 301 | 187 | 153 | 20 | 19 |
| TCGA-OV | 436 | 435 | 173 | 121 | 58 | 47 | 10 | 7 |
| TCGA-PAAD | 178 | 178 | 168 | 50 | 77 | 19 | 24 | 12 |
| TCGA-PCPG | 179 | 179 | 13 | 12 | 5 | 4 | 8 | 5 |
| TCGA-PRAD | 495 | 495 | 59 | 35 | 27 | 15 | 3 | 7 |
| TCGA-READ | 137 | 136 | 475 | 148 | 232 | 70 | 320 | 186 |
| TCGA-SARC | 237 | 237 | 119 | 70 | 45 | 26 | 2 | 0 |
| TCGA-SKCM | 467 | 467 | 841 | 472 | 413 | 229 | 266 | 220 |
| TCGA-STAD | 437 | 437 | 488 | 157 | 211 | 74 | 17 | 14 |
| TCGA-TGCT | 144 | 128 | 23 | 21 | 9 | 8 | 9 | 6 |
| TCGA-THCA | 492 | 492 | 22 | 12 | 6 | 5 | 4 | 3 |
| TCGA-THYM | 123 | 123 | 39 | 24 | 10 | 4 | 6 | 2 |
| TCGA-UCEC | 530 | 530 | 1672 | 149 | 708 | 54 | 118 | 109 |
| TCGA-ACC | 92 | 92 | 117 | 36 | 0 | 0 | 0 | 0 |
| TCGA-CHOL | 51 | 45 | 110 | 62 | 0 | 0 | 0 | 0 |
| TCGA-DLBC | 37 | 37 | 173 | 157 | 0 | 0 | 0 | 0 |
| TCGA-KICH | 66 | 66 | 44 | 25 | 0 | 0 | 0 | 0 |
| TCGA-MESO | 82 | 82 | 47 | 44 | 0 | 0 | 0 | 0 |
| TCGA-UCS | 57 | 57 | 183 | 67 | 0 | 0 | 0 | 0 |
| TCGA-UVM | 80 | 80 | 23 | 16 | 0 | 0 | 0 | 0 |
| Total | 10182 | 10100 | Total number: 3155183 |  | Total number: 1384531 |  | 1055 | 769 |

Despite large differences between variant selection protocols, there is notable overlap between variants deemed recurrent by the above process, and the 470 variants identified in the cancer hotspot datasets [14] (supplementary figure 1). This overlap is strongly dependent on how frequent those variants are observed: there are 54 common variants out of the 61 variants observed more than 10 times over our dataset (> 88%). Among the 819 variants retained for the comparison (see methods for details), only 5 appear among the variants flagged as possible false positive by Chang *et al.* (< 1%).

### Enrichment in known cancer related genes

We observe that recurrent variants occur substantially more frequently in known cancer-related genes than in other genes (figure 1C). Initially approximatively one percent of all observed variants are found in genes that have been described [17], [18] as oncogenes (54 genes) or tumor suppressor genes (71 genes). When recurrent unique protein changes are considered, the fraction of known oncogenes or tumor suppressor genes is substantially increased to 13% and 6.5% respectively. These fractions only marginally increase to 14% and 7% when only the unique protein changes leading to predicted strong binders for frequent HLA-1 types are considered. Supplementary table 4 shows a similar enrichment of known cancer-related genes per cohort. We observe that the enrichment is stronger for oncogenes than for tumor suppressors. This might be expected, as activating mutations in oncogenes are mainly distributed on a few protein positions, while loss of function mutations in tumor suppressors are generally distributed more broadly along the protein sequence.

It is interesting to observe that several of the highly prevalent neo-epitope candidates occur in genes that are involved in known immune escape mechanisms: RAC1:P29S is recurrent in study SKCM (melanoma), is predicted to lead to strong binding neo-epitopes for HLA-A*01:01 and HLA-A*02:01, and is reported to upregulate PD-L1 in melanoma [19]. CTNNB1:S33C is recurrent in studies LIHC (liver hepatocellular carcinoma) and UCEC (uterine corpus endometrial carcinoma), is predicted to lead to strong binding neo-epitopes for HLA-A*02:01, and has been shown to increase the expression of the Wnt-signalling pathway in hepatocellular carcinoma [20], leading to modulation of the immune response [21] and ultimately to tumor immune escape [22]. In a separate study, Cho *et al.* [23] show that this mutation confers acquired resistance to the drug imatinib in metastatic melanoma. Finally, FLT3:D835Y recurrent in study LAML (acute myeloid leukemia), is predicted to lead to a strong binding neo-epitope for HLA-A*01:01, HLA-A*02:01 and HLA-C*06:02, and following Reiter *et al.* [24], Tyrosine Kinase Inhibitors promote the surface expression of the mutated FLT3, enhancing FLT3-directed immunotherapy options, as its surface expression is negatively correlated with proliferation.

While the described mechanisms are probably sufficient to explain immune escape in tumor evolution, the candidates could nevertheless be viable targets for adoptive T cell therapy or TCR gene therapy.

### Recurrent neo-epitopes in patient populations

Upon assumption of statistical independence, the product of the frequency of a recurrent variant with the frequency of class I alleles in the population and the incidence rates of cancer types provides an estimate for the number of patients that carry that specific candidate. Using the number of newly diagnosed patients per year and HLA-1 frequency in the US population, we are able to compute the expected number of patients for 18 cancer entities for which both cancer census data and a TCGA study are available. The occurrence numbers for individual candidates range from 0 to 2,254 for PIK3CA:H1047R in breast cancer patients of type HLA-C*07:01; table 2 presents a summary of expected patient numbers for the complete set of candidates. We estimate that, in the US alone, the previously discussed RAC1:P29S mutation might be present in 628 new patients carrying the HLA-A*02:01 allele each year (in 556 melanoma patients and in 72 lung small cell, head & neck or uterine carcinomas patients, see supplementary table 5 for details). For the CTNNB1:S33C mutation, the total number of HLA-A*02:01 patients in the US is expected to be 364, from uterine corpus, prostate and liver cancer types. As another example, 115 myeloid leukemia patients in the US are expected to be of type HLA-A*02:01 and carry the FLT3:D835Y mutation.

**Table 2:**
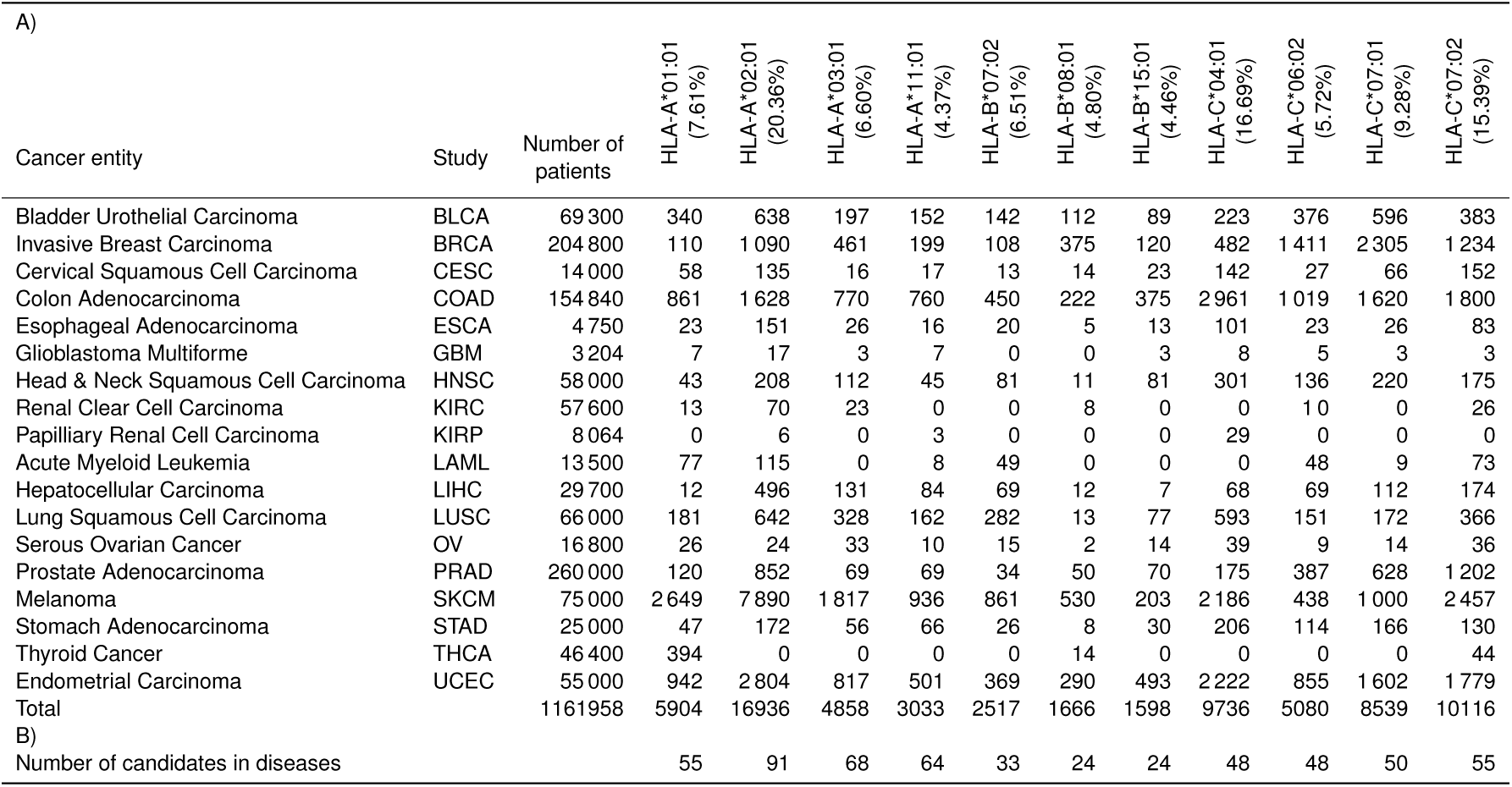
Expected number of newly diagnosed U.S. patients by HLA-1 type and cancer entity. A) Expected number of patients of a given HLA-1 type who harbor at least one potentially immunogenic neo-epitope candidate for that HLA-1 type. Both the cancer incidence and the allele frequency are estimated for the U.S. population. The probability that a patient carries at least one variant from the set of neo-epitope candidates is computed under the assumption that the occurrence of variants in a cancer patient stems from statistically independent events. B) Number of neo-epitope candidates identified in the 18 studies shown in A, which are predicted to be strong binders to the corresponding HLA-1 type.

| A) |  |  |  |  |  |  |  |  |  |  |  |  |  |
| --- | --- | --- | --- | --- | --- | --- | --- | --- | --- | --- | --- | --- | --- |
| Cancer entity | Study | Number of patients | HLA-A*01:01<br>(7.61%) | HLA-A*02:01<br>(20.36%) | HLA-A*03:01<br>(6.60%) | HLA-A*11:01<br>(4.37%) | HLA-B*07:02<br>(6.51%) | HLA-B*08:01<br>(4.80%) | HLA-B*15:01<br>(4.46%) | HLA-C*04:01<br>(16.69%) | HLA-C*06:02<br>(5.72%) | HLA-C*07:01<br>(9.28%) | HLA-C*07:02<br>(15.39%) |
| Bladder Urothelial Carcinoma | BLCA | 69 300 | 340 | 638 | 197 | 152 | 142 | 112 | 89 | 223 | 376 | 596 | 383 |
| Invasive Breast Carcinoma | BRCA | 204 800 | 110 | 1 090 | 461 | 199 | 108 | 375 | 120 | 482 | 1 411 | 2 305 | 1 234 |
| Cervical Squamous Cell Carcinoma | CESC | 14 000 | 58 | 135 | 16 | 17 | 13 | 14 | 23 | 142 | 27 | 66 | 152 |
| Colon Adenocarcinoma | COAD | 154 840 | 861 | 1 628 | 770 | 760 | 450 | 222 | 375 | 2 961 | 1 019 | 1 620 | 1 800 |
| Esophageal Adenocarcinoma | ESCA | 4 750 | 23 | 151 | 26 | 16 | 20 | 5 | 13 | 101 | 23 | 26 | 83 |
| Glioblastoma Multiforme | GBM | 3 204 | 7 | 17 | 3 | 7 | 0 | 0 | 3 | 8 | 5 | 3 | 3 |
| Head & Neck Squamous Cell Carcinoma | HNSC | 58 000 | 43 | 208 | 112 | 45 | 81 | 11 | 81 | 301 | 136 | 220 | 175 |
| Renal Clear Cell Carcinoma | KIRC | 57 600 | 13 | 70 | 23 | 0 | 0 | 8 | 0 | 0 | 10 | 0 | 26 |
| Papillary Renal Cell Carcinoma | KIRP | 8 064 | 0 | 6 | 0 | 3 | 0 | 0 | 0 | 29 | 0 | 0 | 0 |
| Acute Myeloid Leukemia | LAML | 13 500 | 77 | 115 | 0 | 8 | 49 | 0 | 0 | 0 | 48 | 9 | 73 |
| Hepatocellular Carcinoma | LIHC | 29 700 | 12 | 496 | 131 | 84 | 69 | 12 | 7 | 68 | 69 | 112 | 174 |
| Lung Squamous Cell Carcinoma | LUSC | 66 000 | 181 | 642 | 328 | 162 | 282 | 13 | 77 | 593 | 151 | 172 | 366 |
| Serous Ovarian Cancer | OV | 16 800 | 26 | 24 | 33 | 10 | 15 | 2 | 14 | 39 | 9 | 14 | 36 |
| Prostate Adenocarcinoma | PRAD | 260 000 | 120 | 852 | 69 | 69 | 34 | 50 | 70 | 175 | 387 | 628 | 1 202 |
| Melanoma | SKCM | 75 000 | 2 649 | 7 890 | 1 817 | 936 | 861 | 530 | 203 | 2 186 | 438 | 1 000 | 2 457 |
| Stomach Adenocarcinoma | STAD | 25 000 | 47 | 172 | 56 | 66 | 26 | 8 | 30 | 206 | 114 | 166 | 130 |
| Thyroid Cancer | THCA | 46 400 | 394 | 0 | 0 | 0 | 0 | 14 | 0 | 0 | 0 | 0 | 44 |
| Endometrial Carcinoma | UCEC | 55 000 | 942 | 2 804 | 817 | 501 | 369 | 290 | 493 | 2 222 | 855 | 1 602 | 1 779 |
| Total |  | 1 161 958 | 5 904 | 16 936 | 4 858 | 3 033 | 2 517 | 1 666 | 1 598 | 9 736 | 5 080 | 8 539 | 10 116 |
| B) |  |  |  |  |  |  |  |  |  |  |  |  |  |
| Number of candidates in diseases |  |  | 55 | 91 | 68 | 64 | 33 | 24 | 24 | 48 | 48 | 50 | 55 |

Figure 2 shows the cumulative expected number of patients that carry a specific epitope, and with matching HLA-1 type, for the 50 candidates with the highest expected patients number. The number of patients is derived from the sum over all cancer entities, including those in which the candidate is not recurrent according to our criteria. For example, among newly diagnosed US patients of type HLA-C*04:01, 88 prostate cancer patients are expected to carry the mutation PIK3CA:R88Q, even though its observed frequency in the PRAD study is as low as 0.2%. The data shown in figure 3 can be found in supplementary table 5.

**Figure 2:**
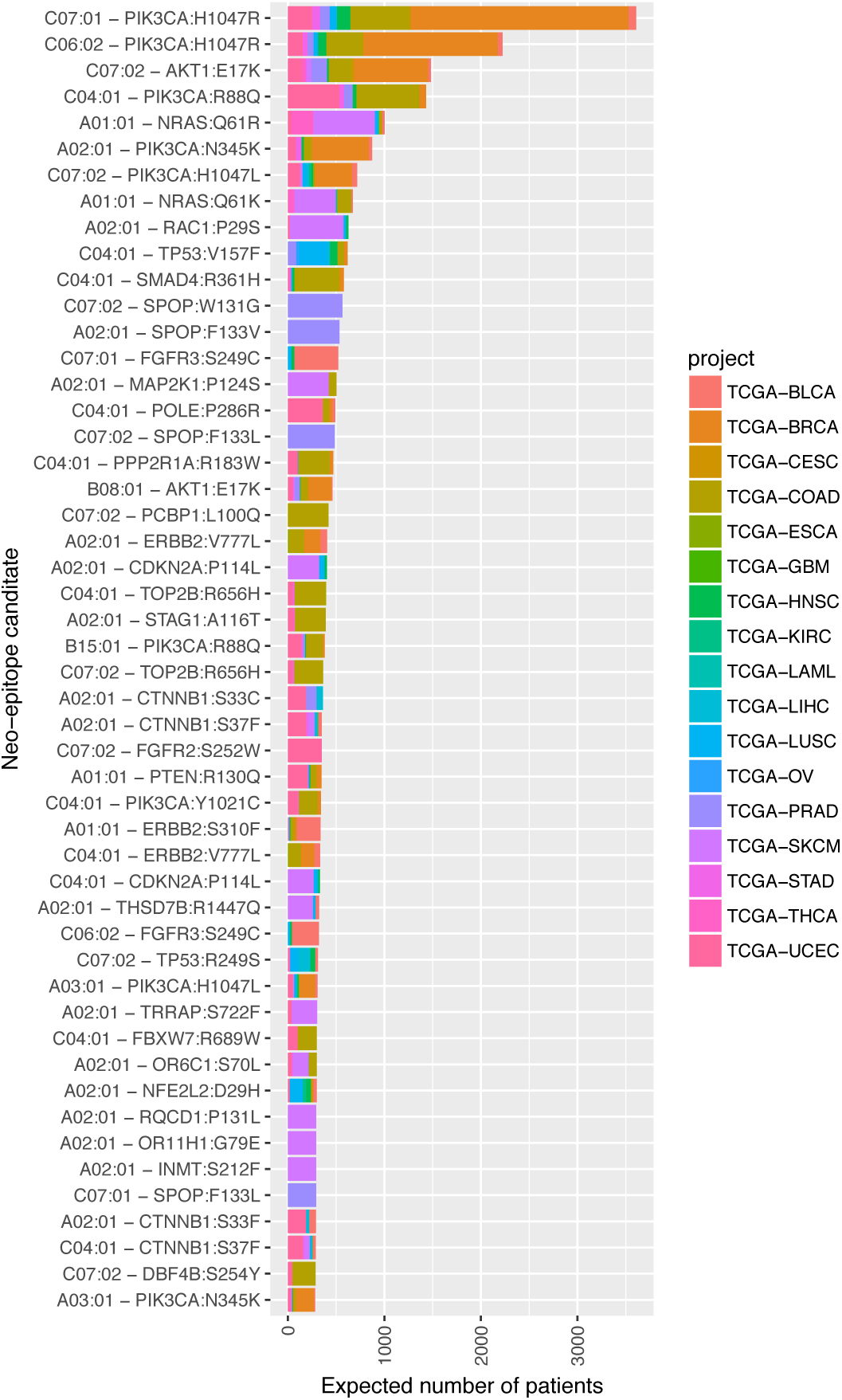
50 most frequent neo-epitope and HLA-1 combinations in patients for which strong MHC I binding is predicted: for each candidate, the expected number of patients is obtained by summing over the 18 cancer entities for which the number of newly diagnosed patients in the US is available, and for which a corresponding TCGA study has been included in our analysis.

**Figure 3:**
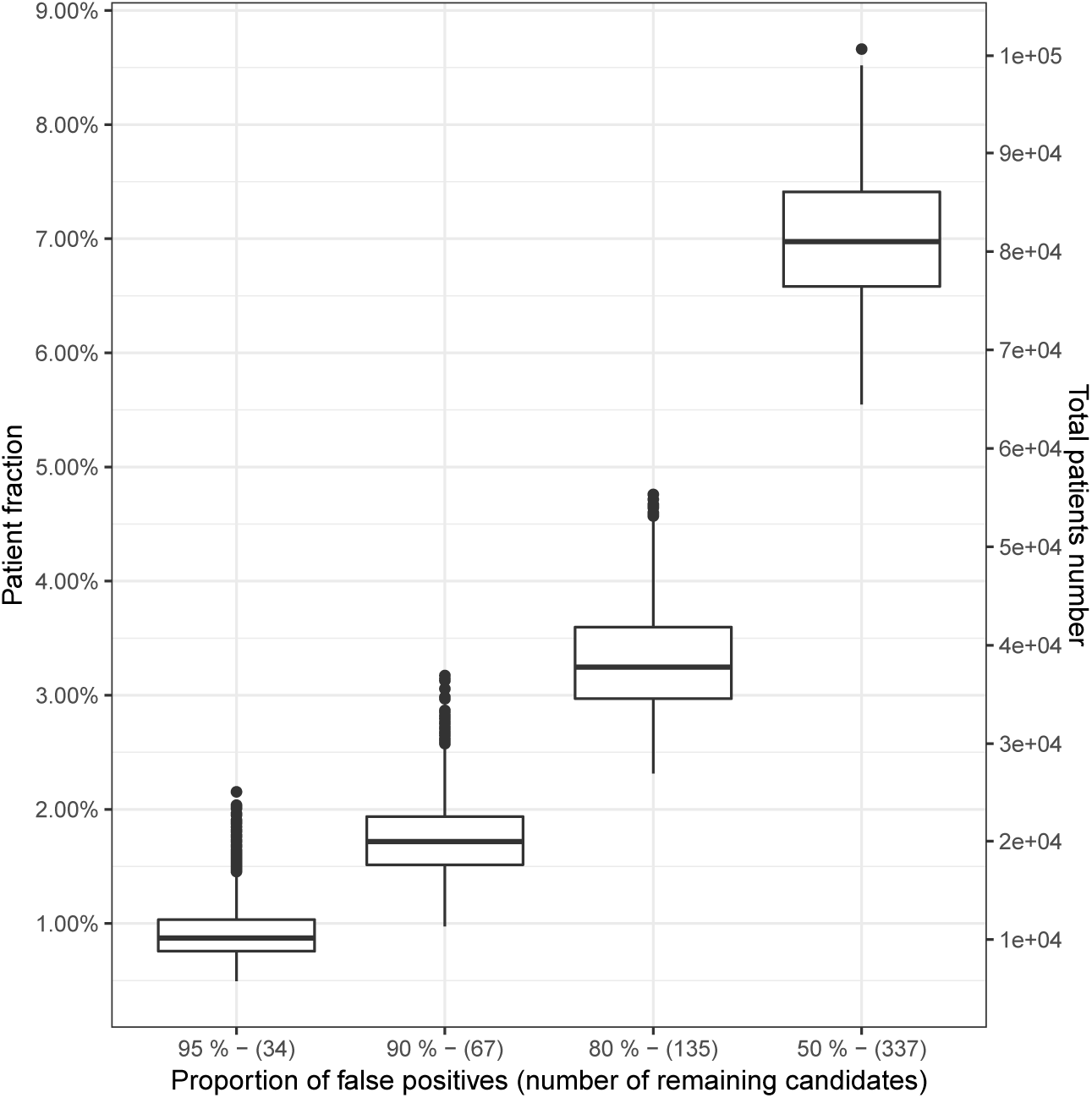
Expected influence of the proportion of false positive neo-epitope candidates on the patient population: Proportion of the patients that carry at least one neo-epitope candidate mutation, and whose HLA-I allele set contains the candidate HLA type, when a limited percentage of the neo-epitope candidates is considered. The patient cohort considered here consists of 6868 patients from the 18 TCGA cohorts for whom the HLA types are known. For each false positive proportion, the false positive candidates have been selected 1000 times at random.

### Candidates libraries

As our current understanding of peptide immunogenicity is still incomplete [25], not all candidates predicted by our pipeline can be expected to trigger an immunogenic response in patients. To estimate the impact of such “false positives”, we have randomly flagged 1000 times 337, 539, 607 & 640 candidates as “false positives”, which is corresponding to a fraction of about 50%, 80%, 90% and 95% of the total 674 candidates. This procedure left us with 1000 sets of 337, 135, 67 & 34 candidates that were not flagged as “false positives”. We have then used a subset of 6868 patients from the 18 TCGA cohorts with cancer census data, for whom HLA types were known. For each candidates subsets, we have counted the number of patients that carry at least one of the remaining candidate mutation, and whose HLA allele set contains the candidate HLA type.

Figure 3 shows that for a pessimistic 90% of false positive candidates, more than 1.5% of patients over all cancer entities (mean 1.78%, median 1.72%, both corresponding to about 20000 new patients per year in the U.S.) are still expected to carry at least one of the 67 remaining candidates’ mutation and corresponding HLA allele. While the proportions are modest, the absolute number of patients Supplementary figure 2 shows that there are considerable differences between entities: the proportion of matching patients is much higher in diseases with high mutational load such as melanomas (TCGA-SKCM, median about 9% for 90% false positives), than in diseases with lower mutational load, such as thyroid cancer (TCGA-THCA, 0.2%, 90% false positives).

### Confirmational evidence

A limited validation of our method was performed in two steps: first, we confirmed that our pipeline was able to identify candidates that have been previously reported as eliciting spontaneous CD8^+^ T-cell responses in cancer patients in whom the target epitopes were subsequently discovered [26], [27]. Both sets together (supplementary table 3) contain 37 epitopes, 35 of which could be mapped to an ENSEMBL transcript (33 unique genes). For 27 of these epitopes our pipeline predicted strong binding with the specific HLA-1 type reported in the corresponding wet-lab investigations. Another 5 epitopes where predicted as weak binders, some of the latter are also predicted to be strong binders in other HLA-1 types. Our pipeline classified 70% of a set of known tumor neo-antigens as strong binders and another 14% as weak binders.

4 out of 34 unique identifiable variants studied by van Buuren *et al.* [27] and Fritsch [26] are found among our set of high confidence missense variants, but only one (CTNNB1:S37F) fulfills the 1% recurrence threshold (9 uterine carcinoma patients). This variant was shown to trigger immunological response against HLA-A*24:02 [28], which isn’t in the set of alleles that we have systematically tested. However, our prediction show that the same peptide might also be reactive against HLAC07:02.

Finally, the CDK4:R24C peptide (sequence ACDPHSGHFV, see supplementary table 3) is not predicted to bind to HLA-A*02:01, even though it leads to confirmed T cell response [29], and has been related to cutaneous malignant melanoma and hereditary cutaneous melanoma [30], [31].

We have also performed preliminary validation for two candidates: RAC1:P29S & TRRAP:S722F binding to HLA-A*02:01 (figure 4). We utilized transgenic mice that harbour the human TCR*αβ* gene loci, a chimeric HLA-A2 gene and are deficient for mouse TCR*αβ* and mouse MHC I genes (termed ABabDII). These mice have been shown to express a diverse human TCR repertoire [32], [33] and thus mimic human T cell response. They were immunized at least twice with mutant peptides and IFN_γ_ producing CD8^+^ T cells were monitored in *ex vivo* ICS analysis 7 days after the last immunization. CD8^+^ T cells were purified from spleen cell cultures of reactive mice using either IFN_γ_-capture or tetramer-guided FACSort. Sequencing of specific TCR *α* and *β* chain amplicons that were obtained by RACE-PCR revealed that this procedure yields an almost monoclonal CD8^+^ T cell population (not shown). In both cases, tested neo-antigen candidates lead to T cell reactivity, confirming not only predicted MHC binding by our pipeline but also immunogenicity *in vivo* in human TCR transgenic mice. Therefore this workflow also allows to generate potentially therapeutic relevant TCRs to be used in the clinics for cancer immunotherapy.

**Figure 4:**
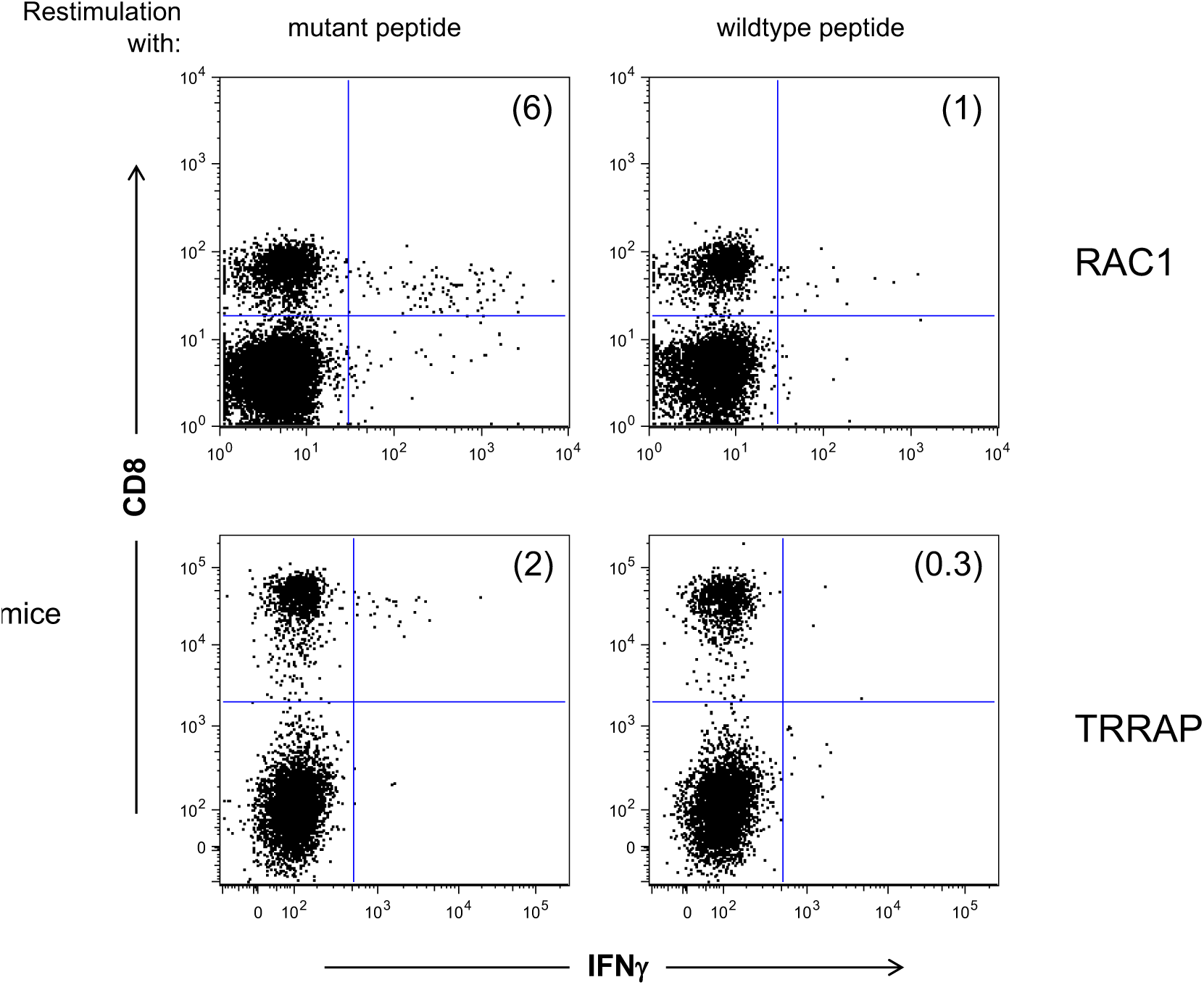
Recognition of predicted epitopes by CD8^+^ T cells: Epitopes for recurrent mutations that have been identified *in silico* to bind to HLA-A*02:01 using our pipeline were synthesized and used for immunization of human TCR transgenic ABabDII mice. Examples (RAC1:P29S and TRRAP:S722F) of *ex vivo* ICS analysis of mutant peptide immunized ABabDII mice 7 days after the last immunization are shown. Polyclonal stimulation with CD3/CD28 dynabeads was used as positive control, stimulation with an irrelevant peptide served as negative control (data not shown).

## Discussion

By virtue of the underlying mutational processes, the genome architecture and accessibility as well as for functional reasons within the disease process, certain somatic mutations will be present in multiple patients while still being highly specific to the tumor [14]. Using existing cancer studies and neo-epitope binding predictions to MHC class I proteins, we propose a ranking of candidates which mutation occur frequently in observed cancer patient cohorts. Taking into account the fact that MHC binding is a necessary but not sufficient condition for T cell activity, and the limitations of MHC binding prediction algorithms, we provide an objective ranking of neo-epitopes based on recurrent variants, as a basis for the development of off-the-shelf immunotherapy treatments.

Despite numerous mechanisms of immune evasion, neo-epitopes are important targets of endogenous immunity [5]. In some cases at least, it has been shown that they contribute to tumor recognition [34], achieve high objective response (in melanoma, see ref. [35], [36]), and a single of them is presumably sufficient for tumor regression [37]. Moreover, positive association has been shown between antigen load and cytolytic activity [38], activated T cells [13] and high levels of the PD-1 ligand [39]. Taken together, these results suggest that neo-epitopes occupy a central role in regulating immune response to cancer, and that this role can be exploited for cancer immunotherapy. Even though the question of negative selection for strong binding neo-epitopes and its relation to other immune evasion mechanisms like HLA loss or PD-L1, CTLA4 dis-regulation is still open [40]. A recent CRISPR screen suggest that more then 500 genes are essential for cancer immunotherapy [41].

Targeting neo-epitopes based on non-recurrent, *private* somatic variants requires generation of private TCRs or CARs for each individual patient, which is challenging [42]. Successful treatments based on genetically engineered lymphocytes has been shown for epitopes arising from unmutated proteins, i.e. *public epitopes*: MART-1 and gp100 proteins have been targeted in melanoma cases [43]. In another trial, Robbins *et al.* [44] have studied long-term follow-up of patients who were treated with TCR-transduced T cells against NY-ESO-1, a protein whose expression is normally restricted to testis, but which is frequently aberrantly expressed in tumor cells. They show that treatment may be effective for some patients. These results show that immune treatments based on *public* variants can be beneficial, suggesting that similar success may potentially be achieved using candidates based on recurrent variants.

However, targeting such non somatic epitopes presents safety and efficacy concerns [2]. The administration of T cells transduced with MART-1 specific T-cell receptor have led to fatal outcomes [45]. Cross-reactivity of TCR against MAGE-A3 (a protein normally restricted to testis and placenta) caused cardiovascular toxicity [46]. Neo-epitopes based on recurrent somatic variants potentially alleviate such problems, as the target sequences are truly restricted to tumor cells.

Our computation of expected targetable patient groups assumes that neither the cancer type nor the patient’s mutanome are associated with the patient’s HLA-1 alleles. In a recent study, Van den Eyden *et al.* [40] show that there is little (if any) antigen depletion due to the negative selection pressure from the immune response. Molecular evolution methods applied to somatic mutations show that nearly all mutations escape negative selection [47]. Taken together, these results suggest that the expected probability of a recurrent variant being present in a patient somatic mutations pool should not be affected (significantly) by the patient’s HLA-1 alleles.

The neo-epitope landscape is diverse and sparse [13]. Few neo-epitopes are predicted to be both strong binders and present in multiple patients. In their analysis, Hartmaier *et al.* [48] estimate that neo-epitopes suitable for precision immunotherapy might be relevant for about 0.3% of the patients, which is in agreement with our results. However, the absolute number of patients is still considerable, see Table 2. Our study shows that a relatively large number of patients (about 1% of newly diagnosed patients) might benefit from a small library of candidates proven to generate immunological response. These numbers must be compared to “conventional” personalised immunotherapy, where a immunologically active candidate must be identified for each new patient for which efficacy and safety are always unknown. Even if a substantial part of the neo-epitopes we suggest turns out to be false positives due to the limitation of prediction algorithms and understanding of immune response, there is potential to help tens of thousands of patients.

## Conclusions

Off the shelf immune treatments can be faster, less costly and safer for individual patients, because each neo-epitope based treatment scheme can be reused on hundreds of patients per year. In this respect, they might open the way to supplement existing personalized cancer immune treatments approaches with precision treatment options.

We believe that our ranking provides a rational order for testing for and selecting off the shelf neo-epitope based therapies. Our preliminary in vivo mouse experiments show that this in principle feasible.

## Materials and Methods

### Data sets

Somatic variants for different cancer entities have been determined using matched pairs of tumor and blood whole exome or whole genome sequencing in the TCGA consortium. We downloaded the open-access somatic variants from GDC data release 7.0 [16], consisting of 33 TCGA projects and 10,182 donors in total. We excluded patients without corresponding entries in the clinical information tables, and 7 projects with less than 100 samples, yielding 9,641 samples covering 26 cancer studies.

### Variant selection

For each sample we selected all single nucleotide variants obtained by the “mutect2” pipeline, that had a “Variant_Type” equal to “SNP”, a valid ENSEMBL transcript ID and a valid protein mutation in “HGVSp_Short”. From these variants, we selected those with a “Variant_Classification” equal to “Missense_Mutation”. We checked that all variants had a “Mutation_Status” equal to (up to capitalisation) “Somatic”, that the total depth “t_depth” was the sum of the reference “t_ref_count” and the alternate “t_alt_count” alleles counts, and that the genomics variant length is one nucleotide. To avoid high number of false positives we consider only variants that are supported by at least 5 reads and have a VAF of at least 10%. Furthermore we removed any variant that occurs with more than 1% in any population contained in the ExAC database version 0.31 [49], by coordinates liftover from the GRCh38 to hg19 human genome versions. This way we obtained 26 cancer entity data sets containing a total of 9,641 samples with an overall 1,384,531 variants.

### Recurrent protein variant selection

We define recurrence strictly on the protein/amino acid exchange level, i.e. different nucleotide acid variants leading to the same amino acid exchange due to code redundancy will be counted together. Recurrent protein variants are defined within each TCGA study. A protein variant is deemed recurrent when it appears in at least 1% of all the patients in the cohort. As cancer types are only considered when the number of patients involved in the studies is greater than 100, this threshold ensures that every recurrent variant has been observed in at least 2 patients for a given cancer type. To be conservative, the recurrence frequency has been computed using, for the denominator, all patients with clinical information in the study, including those without high-confidence missense SNVs. Using this definition, the total number of recurrent amino acid changes is 1,055. A variant recurrent in multiple cancer types is counted multiple times in the above number, the number of unique recurrent variants regardless of the cancer is 869. Supplementary table 1 shows the most frequent amino acid exchanges across 25 cancer entities, as no variant from project TCGA-KIRC’s donors is labeled as recurrent.

Recurrent variants occurring at the same positions (for example when gene’s IDH1 codon R132 is mutated to amino acid H, C, G or S) have been merged into 819 variants suitable for comparisons with the cancer hot spots lists [14]. 122 out of the 819 merged variants belong to the set of 470 cancer hotspot variants, and 5 (PCBP1:L100, SPTLC3:R97, EEF1A1:T432, BCLAF1:E163 & TTN:S3271) to the set of presumptive false positives hotspots listed in the supplementary material of [14].

### Epitopes selection and MHC Class I binding prediction

In the next step all peptide stretches (9-, 10, or 11-mers) containing any of the identified recurrent amino acid exchanges are generated *in silico*. For MHC class I binding prediction we selected 11 frequent HLA-1 types: HLA-A*01:01, HLA-A*02:01, HLA-A*03:01, HLA-A*11:01, HLA-B*07:02, HLA-B*08:01, HLA-B*15:01, HLA-C*04:01, HLA-C*06:02, HLA-C*07:01, HLA-C*07:02. A variety of machine learning algorithms have been developed to determine the MHC binding *in silico*, see ref. [50] for review. Most methods are trained on Immune Epitope Database (IEDB) [51] entries and use allele specific predictors for frequent alleles, while pan-methods are applied to extrapolate to less common alleles. We predicted the MHC class I binding using NetMHCcons [52] v1.1, which predicts peptides IC50 binding, and classifies these predictions as non-binder, weak and strong binders. In the study, we have used these classes to select our neo-epitope candidate.

For a given recurrent variant and a given HLA-1 type, the epitope prediction pipeline can produce multiple overlapping epitopes candidates, differing only by their length. To remove such size redundancy, only the epitope with the lowest predicted mutant sequence IC50 is retained. This procedure also removes non-overlapping epitopes, to keep only at most one epitope per recurrent protein variant, cancer type and HLA-1 type. For comparison we also compute the IC50 for the respective wild type peptide.

This way, we obtain 769 strong binding recurrent peptides and 1829 weak binders. Their complete list is in supplementary table 2, where each candidate is listed with the HLA-1 type it is preticted to bind to.

### Data QC

To ensure that the proportion of variants caused by technical artifacts is small, we have computed the proportion of SNVs called in poly-A, poly-C, poly-G or poly-T repeats of length greater than 6 have been computed for each data study [53], for unique variants (that occur in only one patient across a project cohort), and for variants that are observed more than once in a cohort (supplementary figure 3). For comparison, we have computed the expected frequency of such events, assuming that all possible 11-mers (the mutated nucleotide at the center, flanked by 5 nucleotides on each side) are equiprobable, regardless of their sequence.

Based on this quiprobable model, we have computed the probability that the number of mutations found in repeat locii is equal to or greater than the observed numbers. When considering variants appearing more than once, this probability is not significant for all studies; when unique variants are considered, those appear in repeat locii significantly more often than expected by chance in 7 out of 26 studies (TCGA-COAD, TCGA-KIRP, TCGA-LIHC, TCGA-READ, TCGA-SKCM, TCGA-TGCT & TCGA-UCAC, significance level set to 0.05 after Benjamini-Hochberg multiple testing correction).

### Generation of mutation-specific T cells in ABabDII mice

For immunisation 8-12-week old ABabDII mice were injected subcutaneously with 100 *µ*g of mutant short peptide (9-10mers, JPT) supplemented with 50 *µ*g CpG 1826 (TIB Molbiol), emulsified in incomplete Freund’s adjuvant (Sigma). Repetitive immunizations were performed with the same mixture at least three weeks apart. Mutation-specific CD8^+^ T cells in the peripheral blood of immunized animals were assessed by intracellular cytokine staining (ICS) for IFN_γ_ 7 days after each boost. All animal experiments were performed according to institutional and national guidelines and regulations after approval by the governmental authority (Landesamt für Gesundheit und Soziales, Berlin).

### Patient number estimates and HLA-1 frequencies

HLA-1 frequency data *f*_*h*_ for the U.S. population was retrieved from the Allele Frequency Net Database (AFND) [54]. Frequency data were estimated by averaging the allele frequencies of multiple population datasets from the North American (NAM) geographical region. The major U.S. ethnic groups were included and sampled under the NAM category. Cancer incidence data for the U.S. population (*N*_*d*_) was retrieved from the GLOBOCAN 2012 project of the International Agency for Research on Cancer, WHO [55].

Assuming that the fraction of a recurrent variant in the U.S. population affected by cancer entity *d* (*r*_*d*_) is identical to the observed ratio of that variant in the corresponding TCGA study, the number of patients of HLA-1 type *h* whose tumor contain the variant is expected to be

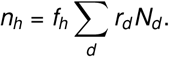

The summation runs over 18 diseases *d* for which both the TCGA projects and the cancer incidence data are available.

## Acknowledgements

Partially supported by Deutsche Forschungsgemeinschaft (SFB-TR36; T.B., G.W.), Deutsche Krebshilfe (111546; G.W., T.B.) and the Berlin Institute of Health (CRG-1; T.B., A.D.). The results presented here are based upon data generated by the TCGA Research Network (http://cancergenome.nih.gov).

## Author Contributions

D.B. conceived and designed the project. E.B. analysed the somatic variant data. E.B., M.H. & C.M. generated neo-epitope candidates, analysed by E.B. G.W. performed *in vivo* validations, and A.D. & E.B. performed the epidemiological analysis. D.B, T.B., G.W. & E.B. contributed to the interpretation of results, and wrote the manuscript.

## Data availability

Somatic variants as well as clinical information were obtained from the GDC repository release 7.0 (https://docs.gdc.cancer.gov/Data/Release_Notes/Data_Release_Notes/#data-release-70). Variant frequencies in the population were downloaded from ExAC (http://exac.broadinstitute.org/), while population HLA-1 allele frequencies were obtained from the AFND database (http://www.allelefrequencies.net/).

## References

1. Schumacher TN, Schreiber RD. Neoantigens in cancer immunotherapy. Science (New York, N.Y.) 2015;348:69–74.

2. Blankenstein T, Leisegang M, Uckert W, Schreiber H. Targeting cancer-specific mutations by T cell receptor gene therapy. Current Opinion in Immunology 2015;33:112–119.

3. Rosenberg SA, Restifo NP. Adoptive cell transfer as personalized immunotherapy for human cancer. Science (New York, N.Y.) 2015;348:62–8.

4. Wirth TC, Kühnel F. Neoantigen Targeting - Dawn of a New Era in Cancer Immunotherapy? Frontiers in Immunology 2017;8:1848.

5. Bethune MT, Joglekar AV. Personalized T cell-mediated cancer immunotherapy: progress and challenges. Current Opinion in Biotechnology 2017;48:142– 152.

6. Kahles A, Lehmann KV, Toussaint NC, Hüser M, Stark SG, Sachsenberg T, Stegle O, Kohlbacher O, Sander C, Cancer Genome Atlas Research Network SJ, et al. Comprehensive Analysis of Alternative Splicing Across Tumors from 8,705 Patients. Cancer cell 2018;34:211–224.e6.

7. Laumont CM, Vincent K, Hesnard L, Audemard É, Bonneil É, Laverdure JP, Gendron P, Courcelles M, Hardy MP, Côté C, et al. Noncoding regions are the main source of targetable tumor-specific antigens. Science translational medicine 2018;10:eaau5516.

8. Liu XS, Mardis ER. Applications of Immunogenomics to Cancer. Cell 2017;168:600– 612.

9. Yarchoan M, Johnson BA, Lutz ER, Laheru DA, Jaffee EM. Targeting neoantigens to augment antitumour immunity. Nature Reviews Cancer 2017;17:209– 222.

10. Luo H, Ye H, Ng HW, Shi L, Tong W, Mendrick DL, Hong H. Machine Learning Methods for Predicting HLA-Peptide Binding Activity. Bioinformatics and biology insights 2015;9:21–9.

11. Gfeller D, Bassani-Sternberg M, Schmidt J, Luescher IF. Current tools for predicting cancer-specific T cell immunity. Oncoimmunology 2016;5:e1177691.

12. Hundal J, Kiwala S, Feng YY, Liu CJ, Govindan R, Chapman WC, Uppaluri R, Swamidass SJ, Griffith OL, Mardis ER, et al. Accounting for proximal variants improves neoantigen prediction. Nature Genetics 2019;51:175–179.

13. Charoentong P, Finotello F, Angelova M, Mayer C, Efremova M, Rieder D, Hackl H, Trajanoski Z. Pan-cancer Immunogenomic Analyses Reveal Genotype-Immunophenotype Relationships and Predictors of Response to Checkpoint Blockade. Cell Reports 2017;18:248–262.

14. Chang MT, Asthana S, Gao SP, Lee BH, Chapman JS, Kandoth C, Gao J, Socci ND, Solit DB, Olshen AB, et al. Identifying recurrent mutations in cancer reveals widespread lineage diversity and mutational specificity. Nature Biotechnology 2016;34:155–163.

15. Wu J, Zhao W, Zhou B, Su Z, Gu X, Zhou Z, Chen S. TSNAdb: A Database for Tumor-specific Neoantigens from Immunogenomics Data Analysis. Genomics, Proteomics & Bioinformatics 2018;16:276–282.

16. Grossman RL, Heath AP, Ferretti V, Varmus HE, Lowy DR, Kibbe WA, Staudt LM. Toward a Shared Vision for Cancer Genomic Data. New England Journal of Medicine 2016;375:1109–1112.

17. Vogelstein B, Papadopoulos N, Velculescu VE, Zhou S, Diaz LA, Kinzler KW, Wunderlich JR, Somerville RP, Hogan K, Hinrichs CS, et al. Cancer genome landscapes. Science (New York, N.Y.) 2013;339:1546–58.

18. Rubio-Perez C, Tamborero D, Schroeder MP, Antolín AA, Deu-Pons J, Perez-Llamas C, Mestres J, Gonzalez-Perez A, Lopez-Bigas N. In Silico Prescription of Anticancer Drugs to Cohorts of 28 Tumor Types Reveals Targeting Opportunities. Cancer Cell 2015;27:382–396.

19. Vu HL, Rosenbaum S, Purwin TJ, Davies MA, Aplin AE. RAC1 P29S regulates PD-L1 expression in melanoma. Pigment Cell & Melanoma Research 2015;28:590–598.

20. Austinat M, Dunsch R, Wittekind C, Tannapfel A, Gebhardt R, Gaunitz F. Correlation between *β*-catenin mutations and expression of Wnt-signaling target genes in hepatocellular carcinoma. Molecular Cancer 2008;7:21.

21. Pai SG, Carneiro BA, Mota JM, Costa R, Leite CA, Barroso-Sousa R, Kaplan JB, Chae YK, Giles FJ. Wnt/beta-catenin pathway: modulating anticancer immune response. Journal of Hematology & Oncology 2017;10:101.

22. Spranger S, Gajewski TF. A new paradigm for tumor immune escape: *β*-catenin-driven immune exclusion. Journal for ImmunoTherapy of Cancer 2015;3:43.

23. Cho J, Kim SY, Kim YJ, Sim MH, Kim ST, Kim NKD, Kim K, Park W, Kim JH, Jang KT, et al. Emergence of CTNNB1 mutation at acquired resistance to KIT inhibitor in metastatic melanoma. Clinical and Translational Oncology 2017;19:1247–1252.

24. Reiter K, Polzer H, Krupka C, Maiser A, Vick B, Rothenberg-Thurley M, Metzeler KH, Dörfel D, Salih HR, Jung G, et al. Tyrosine kinase inhibition increases the cell surface localization of FLT3-ITD and enhances FLT3-directed immunotherapy of acute myeloid leukemia. Leukemia 2018;32:313–322.

25. Gfeller D, Bassani-Sternberg M. Predicting Antigen Presentation-What Could We Learn From a Million Peptides? Frontiers in Immunology 2018;9:1716.

26. Fritsch EF, Rajasagi M, Ott PA, Brusic V, Hacohen N, Wu CJ. HLA-binding properties of tumor neoepitopes in humans. Cancer immunology research 2014;2:522–9.

27. Buuren MM van, Calis JJ, Schumacher TN. High sensitivity of cancer exome-based CD8 T cell neo-antigen identification. OncoImmunology 2014;3:e28836.

28. Robbins PF, El-Gamil M, Li YF, Kawakami Y, Loftus D, Appella E, Rosenberg SA. A mutated beta-catenin gene encodes a melanoma-specific antigen recognized by tumor infiltrating lymphocytes. The Journal of experimental medicine 1996;183:1185–92.

29. Wölfel T, Hauer M, Schneider J, Serrano M, Wölfel C, Klehmann-Hieb E, De Plaen E, Hankeln T, Meyer zum Büschenfelde KH, Beach D. A p16INK4a-insensitive CDK4 mutant targeted by cytolytic T lymphocytes in a human melanoma. Science (New York, N.Y.) 1995;269:1281–4.

30. Landsberg J, Gaffal E, Cron M, Kohlmeyer J, Renn M, Tüting T. Autochthonous primary and metastatic melanomas in Hgf-Cdk4R24C mice evade T-cell-mediated immune surveillance. Pigment Cell & Melanoma Research 2010;23:649–660.

31. Platz A, Ringborg U, Hansson J. Hereditary cutaneous melanoma. Seminars in Cancer Biology 2000;10:319–326.

32. Li LP, Lampert JC, Chen X, Leitao C, Popović J, Müller W, Blankenstein T. Transgenic mice with a diverse human T cell antigen receptor repertoire. Nature Medicine 2010;16:1029–1034.

33. Li L, Blankenstein T. Generation of transgenic mice with megabase-sized human yeast artificial chromosomes by yeast spheroplast-embryonic stem cell fusion. Nature Protocols 2013;8:1567–1582.

34. Robbins PF, Lu YC, El-Gamil M, Li YF, Gross C, Gartner J, Lin JC, Teer JK, Cliften P, Tycksen E, et al. Mining exomic sequencing data to identify mutated antigens recognized by adoptively transferred tumor-reactive T cells. Nature medicine 2013;19:747–52.

35. Rosenberg SA, Dudley ME. Adoptive cell therapy for the treatment of patients with metastatic melanoma. Current Opinion in Immunology 2009;21:233–240.

36. Chandran SS, Somerville RPT, Yang JC, Sherry RM, Klebanoff CA, Goff SL, Wunderlich JR, Danforth DN, Zlott D, Paria BC, et al. Treatment of metastatic uveal melanoma with adoptive transfer of tumour-infiltrating lymphocytes: a single-centre, two-stage, single-arm, phase 2 study. The Lancet. Oncology 2017;18:792–802.

37. Tran E, Turcotte S, Gros A, Robbins PF, Lu YC, Dudley ME, Wunderlich JR, Somerville RP, Hogan K, Hinrichs CS, et al. Cancer immunotherapy based on mutation-specific CD4+ T cells in a patient with epithelial cancer. Science (New York, N.Y.) 2014;344:641–5.

38. Rooney MS, Shukla SA, Wu CJ, Getz G, Hacohen N. Molecular and Genetic Properties of Tumors Associated with Local Immune Cytolytic Activity. Cell 2015;160:48–61.

39. McGranahan N, Furness AJS, Rosenthal R, Ramskov S, Lyngaa R, Saini SK, Jamal-Hanjani M, Wilson GA, Birkbak NJ, Hiley CT, et al. Clonal neoantigens elicit T cell immunoreactivity and sensitivity to immune checkpoint blockade. Science (New York, N.Y.) 2016;351:1463–9.

40. Eynden JV den, Jimenez-Sanchez A, Miller M, Lekholm EL. Lack of detectable neoantigen depletion in the untreated cancer genome. bioRxiv 2018:478263.

41. Patel SJ, Sanjana NE, Kishton RJ, Eidizadeh A, Vodnala SK, Cam M, Gartner JJ, Jia L, Steinberg SM, Yamamoto TN, et al. Identification of essential genes for cancer immunotherapy. Nature 2017;548:537–542.

42. Strønen E, Toebes M, Kelderman S, Buuren MM van, Yang W, Rooij N van, Donia M, Böschen ML, Lund-Johansen F, Olweus J, et al. Targeting of cancer neoantigens with donor-derived T cell receptor repertoires. Science (New York, N.Y.) 2016;352:1337–41.

43. Johnson LA, June CH. Driving gene-engineered T cell immunotherapy of cancer. Cell Research 2017;27:38–58.

44. Robbins PF, Kassim SH, Tran TLN, Crystal JS, Morgan RA, Feldman SA, Yang JC, Dudley ME, Wunderlich JR, Sherry RM, et al. A pilot trial using lymphocytes genetically engineered with an NY-ESO-1-reactive T-cell receptor: long-term follow-up and correlates with response. Clinical cancer research: an official journal of the American Association for Cancer Research 2015;21:1019– 27.

45. Berg JH van den, Gomez-Eerland R, Wiel B van de, Hulshoff L, Broek D van den, Bins A, Tan HL, Harper JV, Hassan NJ, Jakobsen BK, et al. Case Report of a Fatal Serious Adverse Event Upon Administration of T Cells Transduced With a MART-1-specific T-cell Receptor. Molecular Therapy 2015;23:1541– 1550.

46. Linette GP, Stadtmauer EA, Maus MV, Rapoport AP, Levine BL, Emery L, Litzky L, Bagg A, Carreno BM, Cimino PJ, et al. Cardiovascular toxicity and titin cross-reactivity of affinity-enhanced T cells in myeloma and melanoma. Blood 2013;122:863–71.

47. Martincorena I, Raine KM, Gerstung M, Dawson KJ, Haase K, Van Loo P, Davies H, Stratton MR, Campbell PJ. Universal Patterns of Selection in Cancer and Somatic Tissues. Cell 2017;171:1029–1041.e21.

48. Hartmaier RJ, Charo J, Fabrizio D, Goldberg ME, Albacker LA, Pao W, Chmielecki J. Genomic analysis of 63,220 tumors reveals insights into tumor uniqueness and targeted cancer immunotherapy strategies. Genome Medicine 2017;9:16.

49. Lek M, Karczewski KJ, Minikel EV, Samocha KE, Banks E, Fennell T, O’Donnell-Luria AH, Ware JS, Hill AJ, Cummings BB, et al. Analysis of protein-coding genetic variation in 60,706 humans. Nature 2016;536:285–291.

50. Snyder A, Chan TA. Immunogenic peptide discovery in cancer genomes. Current Opinion in Genetics & Development 2015;30:7–16.

51. Vita R, Overton JA, Greenbaum JA, Ponomarenko J, Clark JD, Cantrell JR, Wheeler DK, Gabbard JL, Hix D, Sette A, et al. The immune epitope database (IEDB) 3.0. Nucleic Acids Research 2015;43:D405–D412.

52. Karosiene E, Lundegaard C, Lund O, Nielsen M. NetMHCcons: a consensus method for the major histocompatibility complex class I predictions. Immunogenetics 2012;64:177–186.

53. Laehnemann D, Borkhardt A, McHardy AC. Denoising DNA deep sequencing data-high-throughput sequencing errors and their correction. Briefings in Bioinformatics 2016;17:154–179.

54. González-Galarza FF, Takeshita LY, Santos EJ, Kempson F, Maia MHT, Silva ALSd, Silva ALTe, Ghattaoraya GS, Alfirevic A, Jones AR, et al. Allele frequency net 2015 update: new features for HLA epitopes, KIR and disease and HLA adverse drug reaction associations. Nucleic Acids Research 2015;43:D784– D788.

55. Ferlay J, Soerjomataram I, Dikshit R, Eser S, Mathers C, Rebelo M, Parkin DM, Forman D, Bray F. Cancer incidence and mortality worldwide: Sources, methods and major patterns in GLOBOCAN 2012. International Journal of Cancer 2015;136:E359–E386.

