## Supplemental Data 1 for "Identification and Ranking of Recurrent Neo-Epitopes in Cancer"

### Supplementary tables

**Supplementary table 1:**Table of the 1,055 recurrent variants identified in 26 TCGA studies. For each variant, the number of cases harboring the variant (Number of occurrences), the cohort size and the fraction of cases in the cohort (Fraction) are given. When available, COSMIC entries (from ENSEMBL) are also listed, as well as the highest allele frequency from ExAC version 0.31 (49). Gene annotations from Vogelstein et al. (17) & Rubio-Perez et al. (18) are also provided.

**Supplementary table 2:** Recurrent variants leading to binding (strong and weak binders) neo-epitopes for one of the 11 HLA-1 types considered. Peptide length redundancy has been removed from the variant list, and each variant is listed only once, even if it is recurrent in multiple study cohorts.

**Supplementary table 3:** Confirmation Status with Gold Standard Data Set: the protein changes described in van Buuren et al. (27) and Fritsch et al. (26) have been mapped to the ENSEMBL protein set and neo-epitopes have been computed using our standard pipeline. 26 of these epitopes are exactly recovered by the pipeline, for one of them the pipeline predicts a strong binder for a shorter peptide, and 5 of them are predicted to be weakly binding.

**Supplementary table 4:** Number of tumor suppressor genes and oncogenes from the Vogelstein list (17) by study. The numbers are given for the full set of variants, among recurrent variants only and among neo-epitope candidates. As each protein change is considered only once, the total number of variants is always smaller or equal to the sum over all studies.

**Supplementary table 5:** Neo-epitope candidates for the 18 cancer entities with associated epidemiological data. The candidates are sorted by HLA-1 type, and by protein variant.

### Supplementary figures

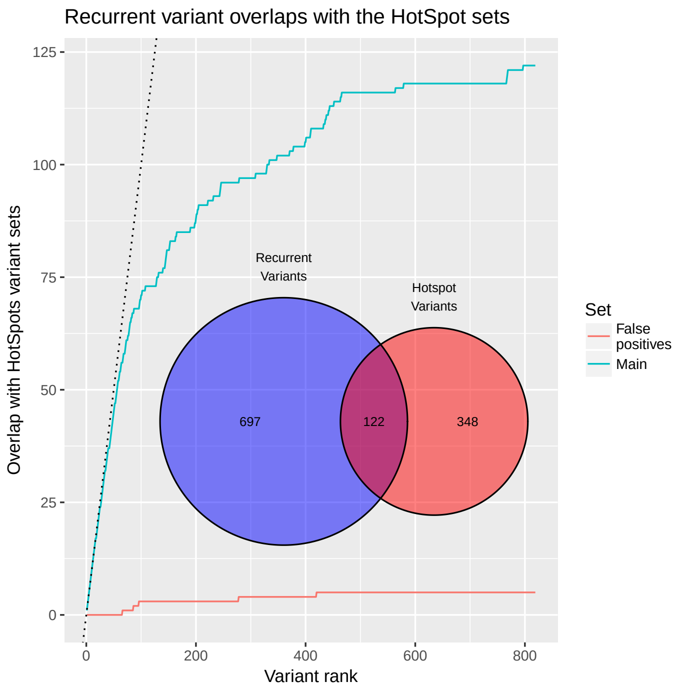


Figure 1: Overlap increase between recurrent variants and Chang *et al.* (14) hotspot variants. The overlap is based on the codon position, so that all variants occurring at the same protein sequence position are pooled together. The recurrent variants that match the alternate codon definition in Chang *et al.* are added to the overlap. The recurrent variants are pooled by codon and sorted by decreasing occurrence frequency in the study. The overlap between hotspots and highly recurrent variants is high, and the common variants fraction decreases when recurrent variants become less frequent. The overlap between recurrent variants and the list of suspected false positive hotspots compiled by Chang *et al.* (14) is very limited. Inset: Venn diagram of the total overlap between the recurrent variants called in this study, and the hotspot variants described in Chang *et al.* (14).

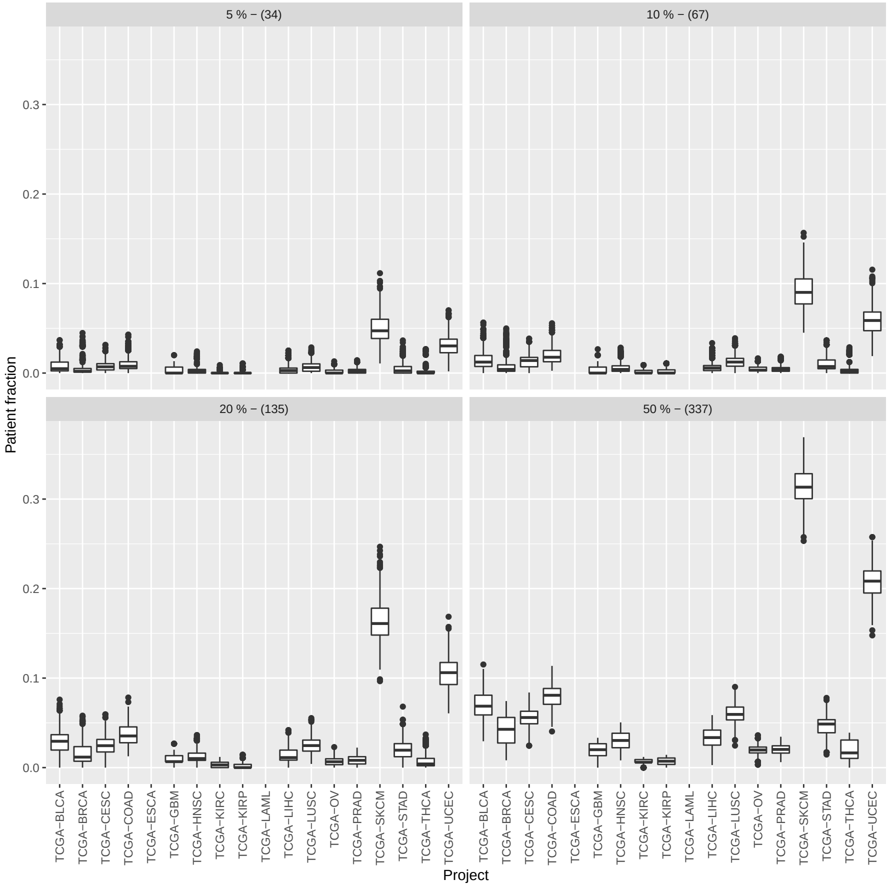


Figure 2: Expected frequency of patients with at least one candidate not labelled as false positive. For each TCGA cohort, we have selected at random 1000 times 50%, 20%, 10% and 5% from the candidates, to conservately model a high rate of false positive within the candidates. From these selected candidates, we have computed the expected frequency of patients with a HLA-1 allele and a mutation matching at least one selected candidate.

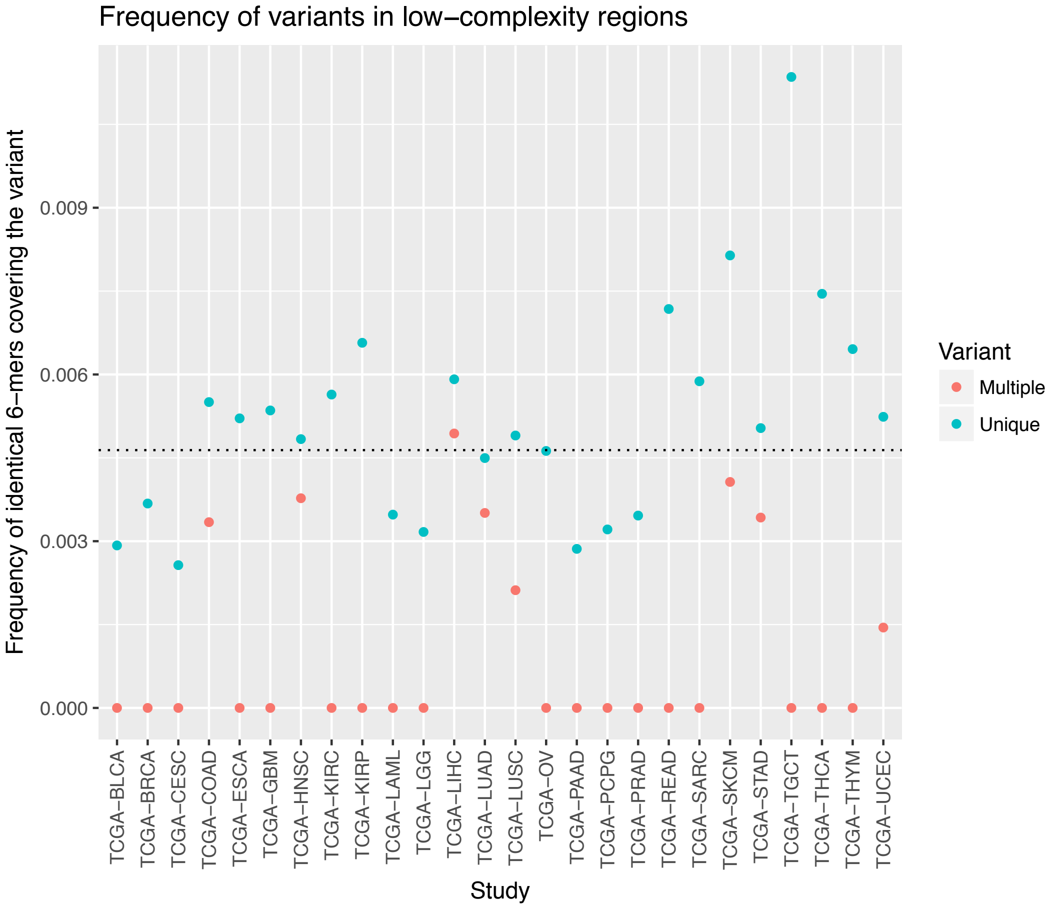


Figure 3: Frequency of Single Nucleotides Variants (SNVs) that fall ina poly-A, poly-C, poly-G or poly-T sequence of length at least 6: the variants that appear only once in the whole study are colored in blue, while the variants that appearmore than once are colored in red. The dotted line shows the expected fraction of suchvariants, if the sequences were all random. Except for the LIHC study, all variants that occur more than once in the cohort are found in difficult-to-sequence regions less oftenthan expected by chance.
